# Real-time integration of cell death mechanisms and proliferation kinetics at the single-cell and population-level using high-throughput live-cell imaging

**DOI:** 10.1101/596239

**Authors:** Jesse D. Gelles, Jarvier N. Mohammed, Luis C. Santos, Diana Legarda, Adrian T. Ting, Jerry E. Chipuk

## Abstract

Quantifying cytostatic and cytotoxic outcomes are integral components of characterizing perturbagens used as research tools and/or in drug discovery pipelines. Furthermore, data-rich acquisition coupled with robust methods for analysis is required to properly assess the function and impact of these perturbagens. Here, we present a detailed and versatile method for Single-cell and Population-level Analyses using Real-time Kinetic Labeling (SPARKL). SPARKL integrates high-content live-cell imaging with automated detection and analysis of fluorescent reporters of cell death. We outline several examples of zero-handling, non-disruptive protocols for detailing cell death mechanisms and proliferation profiles. Additionally, we suggest several methods for mathematically analyzing these data to best utilize the collected kinetic data. Compared to traditional methods of detection and analysis, SPARKL is more sensitive, accurate, and high-throughput while substantially eliminating sample processing and providing richer data.

## Introduction

Programmed cell death pathways are conserved signaling mechanisms which developed early in the evolution of metazoan (Oberst et al., 2008). One aspect shared between many programmed cell death pathways is a variable lag phase between exposure to a perturbagen and the commitment to a cell death program. This lag phase is the consequence of intersecting intracellular pro-death and pro-survival signal transduction and provides a cell with an opportunity to resolve the stress signal and repair accumulated damage (Biton and Ashkenazi, 2011). If these damages are not resolved, the pro-death signaling contributions will overwhelm the pro-survival reserve and trigger biological events committing the cell to death. Importantly, in apoptosis this lag phase also contains an orchestrated and systematic dissolution of organelles and cellular components conducive to efficient clearance with minimal perturbation to neighboring cells. This process is exemplified in apoptosis by the BCL-2 family of proteins, consisting of pro-apoptotic effector proteins (e.g., BAX, BAK) and anti-apoptotic proteins (e.g., BCL-2, BCL-xL), which ultimately serve to regulate the permeabilization of the outer mitochondrial membrane and subsequent activation of the caspase cascade (Wei et al., 2001, Chipuk et al., 2010). However, the kinetics and perpetuation of cell death signaling is highly variable between perturbagens, cell types, death pathways, and between sister cells within a population (Spencer et al., 2009; Gaudet et al., 2012). Elucidating the underlying biology which causes this variability remains a principle focus within the fields of cell death, cell biology, disease etiology, and drug discovery (Kepp et al., 2011). To this end, development of technologies to properly observe and analyze cell death is crucial to progress these fields.

Current standard methods used to observe and quantify cell death remain outdated, suffer from limited throughput, and generate limited datasets for interpretation. The detection and quantification of dead or dying cells is most commonly accomplished by flow cytometry, which requires non-trivial cell numbers, extensive sample handling, sample exposure to significant mechanical and chemical stress, and creates considerable delays between sample harvesting and analyses (Koopman et al., 1994). For example, experiments must be terminated in order to be analyzed, and therefore only provide static endpoint data, requiring considerable effort to optimize the experimental design.

Commonly used reagents involve cell-impermeable “viability” dyes (such as propidium iodide, DRAQ7, SYTOX, and YOYO3), which label cells following loss of plasma membrane integrity and permeabilization. Reliance on this feature for quantification does not distinguish between distinct pathways and labels cells at the tail-end of the dismantling process, thereby failing to capture the time in which cells undergo key biological processes (Vanden Berghe et al., 2010; Dillon et al., 2014). Additionally, labeling with viability dyes is not stoichiometric and often results in pseudo-binary labeling profiles following the first instance of membrane instability. Enzymatically-cleaved fluorescently-conjugated probes (e.g., DEVD-containing caspase-target peptides) are another common strategy despite their cost, difficulty of use, poor signal, and differential and non-specific activation (Yu et al., 2001; McStay et al., 2014). Alternative methodologies use metabolic activity or biochemical measures as surrogate readouts for cell viability, but interpretations from this data are obfuscated by the underlying biology of the perturbagens and ultimately do not directly rely on cell death machinery (Chan et al., 2013).

### Design

Here, we integrate and advance multiple previously described methods for observing and quantifying cell proliferation and cell death using live-cell high-content imagers and refer to this workflow as Single-cell and Population-level Analyses using Real-time Kinetic Labeling (SPARKL). We utilize fluorescently-labeled Annexin V due to stability, signal strength and longevity, stoichiometric binding, and relevance to cell death machinery. Additionally, we test and verify several fluorescent labeling reagents compatible with long-term incubation and time-course studies as well as providing examples of how they can be used to investigate novel aspect of cell death kinetics which were not feasible with analogous workflows. Our non-toxic label-and-go methods outperform previous methodologies while maintaining high-sensitivity, high-accuracy, and a high-throughput zero-handling rapid protocol. Furthermore, we expand on the versatility of our method by providing several example adaptations and mathematical analyses to explore the depth of the collected kinetic data. Collectively, we will demonstrate how data gathered and analyzed using SPARKL can be used to investigate fundamental aspects of cell death biology by applying these workflows to models of death receptor-mediated apoptosis, the mitochondrial pathway of apoptosis, ferroptosis, and necroptosis.

## Results and Discussion

### Automated live-cell imaging and detection of fluorescent labeling provides rich datasets for analyzing, characterizing, and identifying cell death in response to perturbagens

We developed SPARKL chiefly to capture kinetic cell death data since traditional methods utilizing flow cytometry are limited to endpoint data. Additionally, we aimed to design a workflow compatible with high-throughput studies while requiring minimal handling and retaining high sensitivity and specificity. Here, we capitalize on the advent of high-content in-incubator live-cell imagers to capture cellular phenotypes, measure fluorescent signals, and analyze data in real-time through regular scanning schedules. This technology eliminates the requirement for disruptive handling (e.g., trypsinization, pipet-induced shear forces, centrifugation, incubation in labeling buffers, etc.) and is non-invasive to cells in culture. The repeated observation of samples and coupled automated real-time analysis also collects richer datasets in a workflow that is less labor- and time-intensive than analogous workflows using flow cytometry (Figure 1A).

**Figure 1.**
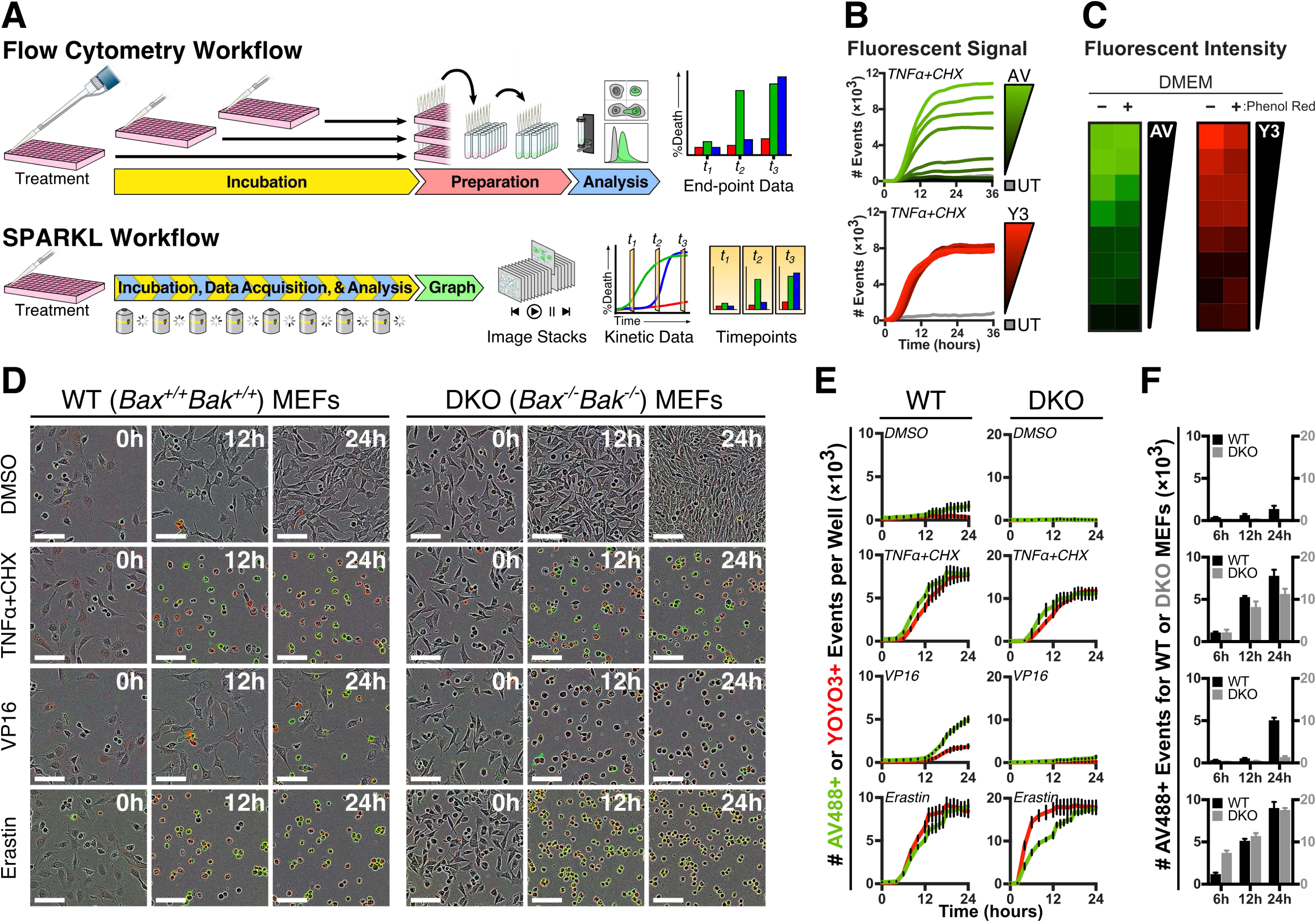
SPARKL provides data-rich kinetic analyses of cell death in real-time using live-cell imaging. (**A**) Comparison of workflows for cell death detection assays using traditional flow cytometry methods or SPARKL live-cell imaging. SPARKL eliminates sample preparation and handling while coupling real-time detection and analysis into a simple, rapid, and less laborious protocol. (**B**) MEFs were co-treated with CHX (50 μg/ml) and mTNFα (10 ng/ml) to induce cell death, incubated with titrations of Annexin V-FITC (67, 100, 133, 200, 400, 1000, 2000, and 4000 ng/ml) and YOYO3 (200, 250, 333, 400, 500, 667, 1000, and 2000 nM), and subjected to the SPARKL protocol. (**C**) MEFs cultured in DMEM lacking or containing phenol red were treated as in ***B***. Positive labeling events were analyzed for signal intensity as mean fluorescence intensity integrated over the event area. Relative fluorescence intensity is depicted as a heat map for each titration of Annexin V-FITC and YOYO3. (**D**) *Bak^+/+^Bax^+/+^* (WT) and *Bak^−/−^Bax^−/−^* (DKO) MEFs were treated with DMSO (0.2%), mTNFα (10 ng/ml) + CHX (50 μg/ml), VP16 (25 μM), or Erastin (10 μM), and incubated with Annexin V-FITC (250 ng/ml) and YOYO3 (250 nM). Images were collected by an IncuCyte ZOOM with a 10× objective and the shown panels are representative images from the stack; 100 μm scale bar. Images were white-point corrected to normalize visualization of green and red pixel background in images lacking a significant fluorescent signal. (**E**) Kinetic data analyzed from images stacks corresponding to cells from ***D***. Dissimilar range of axis reflects differences in number of total cells due to rates of proliferation. Axis range is determined by internal controls within each experiment. (**F**) Comparative endpoint data extracted from ***E*** at the indicated time-points.

One significant benefit of this method is that cells are incubated in normal growth media containing fluorescent probes in order to eliminate stepwise processing to label samples. Therefore, detection reagents must be non-disruptive, highly specific, culture-stable, non-labile, and readily detectable when incubated with cells for prolonged periods of time. Most fluorescent probes utilized in flow cytometry assays depend on the brief incubation period and single-scan detection for their utility and are not compatible in kinetic assays due to inherent cyto-static/-toxic effects with prolonged incubation and rapid photobleaching with successive detection. Therefore, we sought to find reagents which are compatible with kinetic assays by demonstrating no negative effects on cells in culture while exhibiting potent and stable fluorescent signal. Annexin V labels apoptotic cells by detecting phosphatidylserine (PS) on the outer leaflet of the plasma membrane, which is exposed during caspase-dependent cell death (Devaux, 1991). We began these studies by testing FITC-labeled recombinant Annexin V and cell-impermeable viability dye YOYO3 for compatibility within our method (Logue et al., 2009; Gelles and Chipuk, 2016). MEFs were cultured in growth media containing Annexin V and YOYO3, co-treated with tumor necrosis factor α (TNFα) and cycloheximide (CHX) to promote apoptosis, and subjected to SPARKL using a high-content in-incubator live-cell imager. The kinetic data captured the lag phase followed by the robust labeling of MEFs undergoing death (Figure 1B). Here, cells were cultured in media containing sufficient calcium for robust Annexin V labeling, but alternative media formulations may require calcium supplementation (Meers and Mealy, 1993). Importantly, cells not treated with TNFα and CHX exhibited no toxicity as a consequence of either fluorescent probe at the highest concentrations (Figure 1B, “UT” line). As expected, the stoichiometric method of Annexin V-labeling was observed as the concentration-dependent decrease in number of positive events. Said another way, low concentrations of Annexin V will bind to cells exposing PS but may not generate sufficient signal (as defined by the intensity, size, and segmentation of the fluorescent event) according to the user-defined processing definition (see Materials and Methods). However, both Annexin V and YOYO3 were capable of strong and sustained labeling in live-cell imagers at concentrations approximately 10-fold below what is needed in flow cytometric assays (Gelles and Chipuk, 2016). To investigate if growth media containing colorimetric indicators might affect detection of fluorescent signals, we compared signal detection of MEFs instigated to die when incubated in DMEM lacking or containing phenol red. Events of cells labeled with either Annexin V-FITC or YOYO3 exhibited diminished average fluorescent intensity when culture media contained phenol red (Figure 1C). Therefore, all future applications of SPARKL utilized phenol red-free culture media.

The foundation of this method is the collection of high-content images of cells in culture at regular intervals which provide phenotypic characterization of cell populations, confluency data, and detection of fluorescent pixels. To demonstrate the diversity and specificity of data collected with SPARKL, *Bax^+/+^Bak^+/+^* and *Bax^−/−^Bak^−/−^* (WT and DKO, respectively) MEFs were cultured in media containing Annexin V-FITC and YOYO3, treated with perturbagens, and subjected to the SPARKL workflow. Images from select timepoints were manually reviewed to observe cellular morphology and detection of fluorescent labeling (Figure 1D). Green and red pixels from each collected image were quantified using a trained algorithm and reported as the number of Annexin V- and YOYO3-positive events observed at each timepoint (Figure 1E). To assess the selectivity our fluorescent probes, MEFs were instigated to die through the extrinsic pathway of apoptosis (via co-treatment of TNFα+CHX), the intrinsic pathway of apoptosis (via topoisomerase inhibitor, VP16), or non-apoptotic pathway of ferroptosis (via cysteine-glutamate antiport inhibitor, erastin) (Peltzer et al., 2016; Dixon et al., 2012). Both Annexin V and YOYO3 demonstrated accuracy and selectivity by labeling cells in a time-dependent manner. Interestingly, VP16-treated MEFs exhibited the longest lag phase before onset of death and demonstrated Annexin V-positivity more rapidly than YOYO3-positivity, which reflects the biology of PS exposure prior to loss of plasma membrane integrity during apoptosis. To directly compare Annexin V-positivity between WT and DKO MEFs, data collected at specific timepoints were extracted and visualized side-by-side (Figure 1F). It is worth noting that WT and DKO MEFs proliferate at different rates and therefore result in vastly different populations over the course of a short experiment (exampled in Figure 1D). As such, the y-axes for the two cell lines are not congruent but are graphed appropriately based on internal controls (see Materials and Methods); we explore methods for appropriate data normalization in a later section. An important aspect of this methodology is that all data are collected via cell imaging which means all data generated via SPARKL can be backtracked to the original visualization of the cells in culture. As an example, collection and visualization of these data from DKO MEFs treated with VP16 reveals that the morphological phenotypes observed in these cells does not coincide with probe labeling and therefore is not indicative of cell death (Figures 1D-F). In this way, we have demonstrated the depth of data collected using SPARKL and how it can be visualized for thorough interpretation of experimental results.

Investigations of apoptotic cell death commonly employ Annexin V-binding assays due to the exposure of PS on the outer leaflet of the plasma membrane following caspase-mediated cleavage of flippases (Segawa et al., 2014). Exposure of PS is indicative of the underlying apoptotic signaling, and precedes loss of plasma membrane integrity caused by breakdown of cellular homeostatic processes. Therefore, apoptotic studies occasionally utilize a dual-labeling methodology of Annexin V with a cell-impermeable viability dye (such as propidium iodide or DRAQ7) in order to characterize cells in early- or late-apoptosis by dichotomizing single-positive and dual-positive events (Jiang et al., 2016). However, cells undergoing non-apoptotic cell death often do not exhibit this sequential labeling because loss of plasma membrane integrity occurs without the preceding exposure of PS. As a consequence, cells label with the viability dye and Annexin V simultaneously, which is indistinguishable from cells in late-apoptosis when analyzed by flow cytometry (Figure 2A). Therefore, we investigated whether the kinetic data of SPARKL could distinguish between apoptotic and non-apoptotic cell death following treatment of cellular perturbagens.

**Figure 2.**
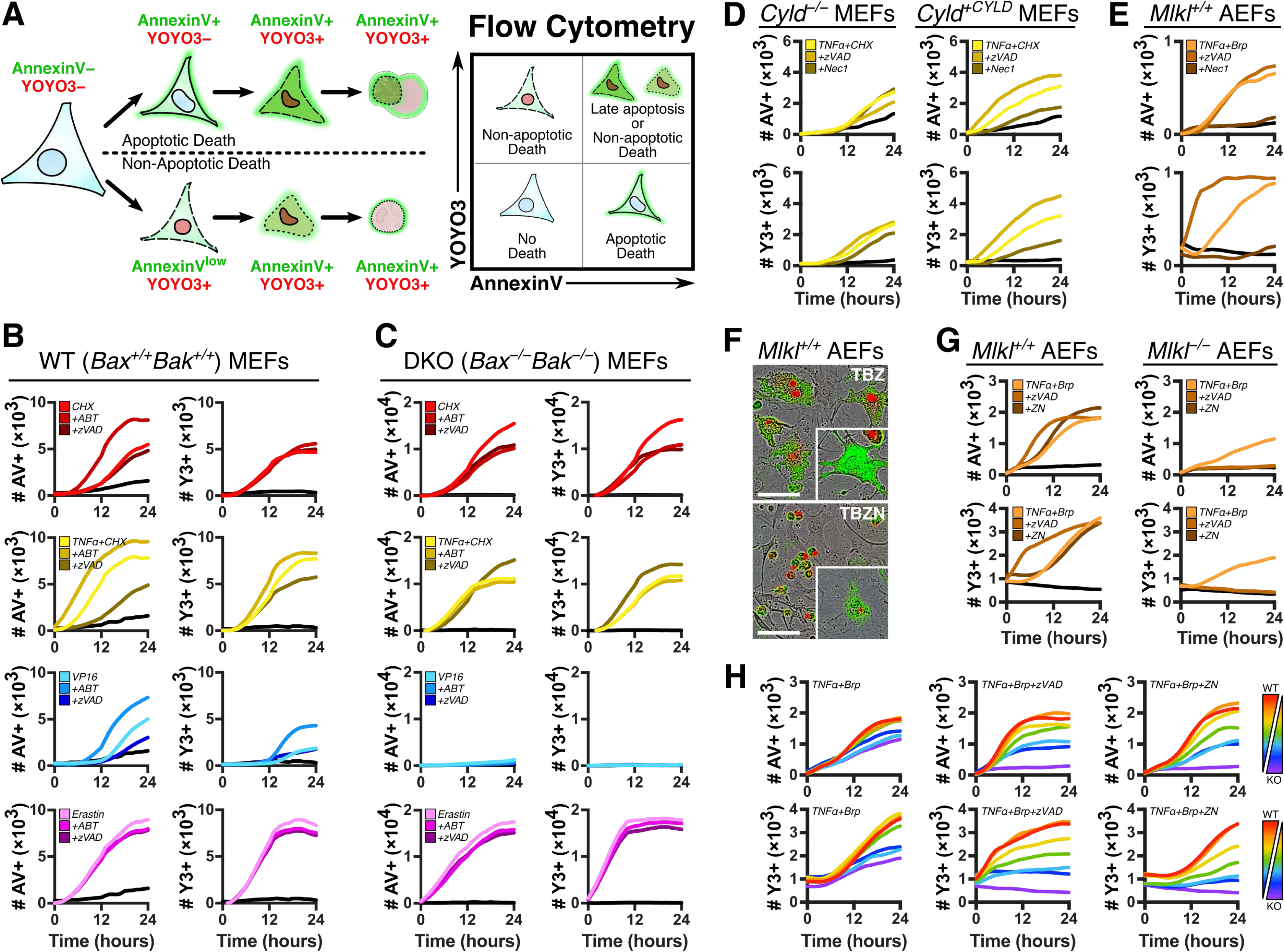
SPARKL is versatile and capable at quantifying and interrogating different pathways of cell death. (**A**) Cells dying by apoptotic or non-apoptotic mechanisms exhibit differential labeling of Annexin V or a cell viability dye (YOYO3) which are not clearly differentiated by flow cytometric methodologies. (**B**) WT MEFs were incubated in growth media containing Annexin V-FITC (250 ng/ml) and YOYO3 (250 nM), treated with CHX (50 μg/ml), mTNFα (10 ng/ml) + CHX (50 μg/ml), VP16 (25 μM), or Erastin (10 μM), co-treated with DMSO, ABT-737 (1 μM), or zVAD-fmk (100 μM), and subjected to SPARKL. (**C**) DKO MEFs were treated and incubated as in ***B***. (**D**) *Cyld^-/-^* MEFs reconstituted with either GST or CYLD were incubated in growth media containing Annexin V-FITC (250 ng/ml) and YOYO3 (250 nM), mTNFα (20 ng/ml) + CHX (50 μg/ml), zVAD-fmk (40 µM), and necrostatin-1 (10 µM), and subjected to SPARKL. (**E**) Primary adult ear fibroblasts (AEFs) were cultured from *Mlkl^+/+^* and incubated in growth media containing Annexin V-FITC (250 ng/ml) and YOYO3 (250 nM), mTNFα (50 ng/ml) + Birinapant (5 µM), zVAD-fmk (40 µM), and necrostatin-1 (10 µM), and subjected to SPARKL. (**F**) Representatives images collected by the IncuCyte ZOOM from *Mlkl^+/+^* AEFs treated with mTNFα (50 ng/ml), Birinapant (5 µM), zVAD-fmk (40 µM), with or without necrostatin-1 (10 µM). Cells are labeled with Annexin-FITC and YOYO3 (green and red, respectively). Stitched insets are a conserved magnification. Scale bars represent 100 µm. (**G**) AEFs from either *Mlkl^+/+^* or *Mlkl^-/-^* mice were treated with mTNFα (50 ng/ml) + Birinapant (10 µM), and co-treated with zVAD-fmk (40 µM), with or without necrostatin-1 (10 µM), and subjected to SPARKL in the presence of Annexin V-FITC (250 ng/ml) and YOYO3 (250 nM). (**H**) WT and *Mlkl* KO AEFs were cocultured in a variety of ratios and treated as in ***G***.

In order to assess the capability of SPARKL to dichotomize kinetic labeling patterns of cells undergoing apoptotic or non-apoptotic cell death, we cultured WT MEFs in growth media containing Annexin V-FITC, YOYO3, and perturbagens engaging either the extrinsic pathway of apoptosis (TNFα+CHX), intrinsic pathway of apoptosis (CHX or VP16), or non-apoptotic ferroptosis (erastin) (Figure 2B). Cell populations instigated to die by TNFα+CHX co-treatment or VP16 exhibited Annexin V-positivity prior to YOYO3-positivity, indicative of apoptotic cell death. By contrast, cells instigated to die by high concentrations of CHX alone or erastin demonstrated similar kinetics of labeling with Annexin V and YOYO3. Additionally, we co-treated cells with ABT-737 (an inhibitor to multiple anti-apoptotic BCL-2 proteins) and/or zVAD-fmk (a pan-caspase inhibitor) to further characterize mechanisms of cell death in response to each perturbagen using the SPARKL assay (Oltersdorf et al., 2005; McStay et al., 2008). Cells dying via apoptosis demonstrated accelerated or attenuated rates of Annexin V-labeling in response to co-treatment with ABT-737 or zVAD-fmk, respectively; cells dying via ferroptosis demonstrated no change in kinetics of Annexin V labeling. By comparison, kinetics of YOYO3 labeling exhibited less resolution and only modest changes in cells co-treated with ABT-737 or zVAD-fmk. Death receptor (DR)-mediated apoptosis can occur either through a mitochondrial independent or dependent pathway, termed Type I and Type II respectively, and the latter signals through members of the BCL-2 family of proteins (Jost et al., 2009). Kinetic data collected by the SPARKL workflow clearly revealed that WT MEFs treated with TNFα+CHX signal primarily via the Type II pathway as evidenced by the synergic labeling of Annexin V in cells co-treated with ABT-737. To further interrogate the mechanism of cell death engaged by each perturbagen, we replicated this treatment strategy in DKO MEFs (Figure 2C). DKO MEFs treated with VP16 demonstrated minimal labeling due to the requirement of BAX and BAK for mitochondrial outer membrane permeabilization in the intrinsic pathway of apoptosis. By contrast, DKO MEFs treated with erastin exhibited similar kinetics of death compared to WT MEFs. DKO MEFs treated with TNFα+CHX died with similar kinetics as their WT counterparts consistent with death receptor signaling which is independent of BAX and BAK. Furthermore, DKO MEFs exhibited similar kinetics for both Annexin V and YOYO3. Interestingly, DKO MEFs treated with CHX alone exhibited death similar to analogously treated WT MEFs but did not synergize with ABT-737, suggesting that CHX treatment engaged multiple pathways of cell death which were selectively amplified with particular co-treatments or in different model systems.

Until this point, we have generalized programmed cell death pathways as either apoptotic or non-apoptotic, the latter of which has been modeled exclusively by ferroptosis thus far. However, death receptor signaling can initiate non-apoptotic cell death programs, such as necroptosis, which are instigated when caspases are inhibited (Vercammen et al., 1998; Kawahara et al., 1998). Interestingly, while WT MEFs treated with TNFα+CHX demonstrated attenuated cell death when co-treated with zVAD-fmk, suggestive of apoptotic cell death, similarly treated DKO MEFs exhibited robust death that modestly increased with co-treatment of zVAD-fmk suggestive of necroptotic cell death (Figure 2B-C). When caspases are inhibited, the switch from DR-mediated apoptosis to necroptosis is initiated by a RIPK1-RIPK3 interaction, which is regulated by the ubiquitylation profile of RIPK1, resulting in RIPK3-mediated phosphorylation of MLKL, which consequentially oligomerizes and permeabilizes the plasma membrane (Sun et al., 2012). It has been reported that SV40-transformed MEFs do not express RIPK3 sufficiently to induce a strong necroptotic program and so we expanded our investigations to different established cellular models (Moujalled et al., 2013). First, we utilized a previously described model for studying necroptosis in which MEFs deficient in CYLD, a RIPK1 deubiquintinase, were reconstituted with either CYLD or a vehicle control (O’Donnell et al., 2011). *Cyld*^−/−^ MEFs treated with TNFα+CHX exhibited cell death but only the MEFs with reconstituted CYLD demonstrated increased or decreased death following co-treatment with zVAD-fmk or the RIPK1 inhibitor, Necrostatin-1 (Nec1), respectively, indicative of necroptosis (Degterev et al., 2005) (Figure 2D). Additionally, MEFs dying by necroptosis exhibited similar labeling kinetics of Annexin V and YOYO3, which parallels the labeling phenotype we observed in MEFs dying by ferroptosis.

Thus far, we relied on CHX co-treatment to reveal pro-death pathways by inhibiting translation of induced pro-survival genes downstream of TNFα signaling. In order to better study labeling kinetics of cells dying by necroptosis we utilized a SMAC mimetic, birinapant (Brp), to inhibit RIPK1 ubiquitylation by cIAP1/2, which facilitates the pro-survival signal cascade (Holler et al., 2000). As a consequence of inhibiting RIPK1 unbiquitylation, the requirement of CYLD to initiate necroptosis was greatly reduced and therefore we moved to an MLKL knockout model for our necroptosis investigations (data not shown). Primary *Mlkl^+/+^* adult ear fibroblasts (WT AEFs) were cultured from mice and treated with TNFα+Brp with- or without co-treatment of zVAD-fmk or Nec1. While Annexin V labeling kinetics appeared unchanged in WT AEFs instigated to undergo necroptosis by addition of zVAD-fmk, YOYO3 demonstrated a dramatic and rapid labeling profile specific to a necroptotic outcome (Figure 2E). Necroptosis is defined by the regulated permeabilization of the plasma membrane by oligomeric MLKL and therefore the rapid labeling by a viability dye is consistent with this phenotype (Wang et al., 2014). However, the unchanged labeling kinetics of Annexin V was unexpected, particularly for cells which became necroptotic hours prior. Inspection of the images collected during SPARKL revealed a substantially distinct cellular morphology in necroptotic cells in which WT AEFs did not contract, retained a topologically distinct and normal nucleus, and slowly exposed PS resulting in a weak Annexin V signal over the large area of the cell (Figure 2F, upper panel). Curiously, a subpopulation of the cells exhibited the same weak Annexin V signal but demonstrated no YOYO3 labeling (Figure 2F, upper inset panel). It has been observed that necroptotic cells are capable of exposing PS following MLKL activation but prior to complete loss of plasma membrane integrity which could explain our observations that WT AEFs slowly accumulate Annexin V labeling following induction of necroptosis (Gong et al., 2017). Labeled WT AEFs co-treated with Nec1 resembled an apoptotic phenotype characterized by cellular contraction and formation of apoptotic bodies; however, there remained a few cells in the population which resembled necroptosis (Figure 2F, bottom panel). To verify that our observations were indicative of necroptosis, we repeated the experiment using *Mlkl^−/−^* primary adult ear fibroblasts (KO AEFs) and observed that these cells were both less sensitive to TNFα+Brp and also completely protected by addition of zVAD-fmk, suggestive of apoptotic cell death (Figure 2G). Finally, we treated cocultured WT and KO AEFs with necroptotic inducers and analyzed the cells with SPARKL to determine if the pattern of labeling would reflect the heterogeneity of the population. As the ratio of WT to KO AEFs decreased, we observed a concomitant decrease or absence of TNFα+Brp sensitivity and necroptotic labeling signatures (Figure 2H).

It is worth noting that minimal cell death was observed in TNFα+Brp treated WT AEFs when co-treated with Nec1 (Figure 2E). Paradoxically, co-treating WT AEFs with both zVAD-fmk and Nec1 (TBZN) did not abolish cell death but instead resembled the TNFα+Brp only condition and exhibited apoptosis-like morphology (Figure 2F-H). One explanation is that cells unable to undergo necroptosis eventually resolve the zVAD-induced inhibition of caspases and revert to an apoptotic program. Alternatively, they may be undergoing a necroptotic program that either does not require, or is revealed by the inhibition of, RIPK1 (Lin et al., 2016; Newton et al., 2016). The KO AEFs do not parallel these observations and while this could suggest that death in response to TBZN is MLKL-dependent, data collected with SPARKL revealed that TBZN-induced cell death does not resemble necroptosis by either labeling kinetics or morphology. Nec1 binds the kinase domain of RIPK1, which is responsible for propagating the pro-death signal cascade, and has been shown to selectively inhibit necroptosis in cells induced via a death receptor and co-treatment with CHX±zVAD-fmk (Degterev et al., 2008). However, both apoptosis and necroptosis are inhibited by Nec1 in cells which are induced via a co-treatment with a SMAC mimetic, due to requirement of RIPK1 for the formation of the caspase-8 activating complex, and we observed this differential effect of Nec1 in our studies as well (Wang et al., 2008) (Figures 2D-E).

We validated that the SPARKL workflow is not only high-throughput and less labor intensive, but also versatile and capable of observing and characterizing cell death following exposure to perturbagens. Specifically, we demonstrated that kinetic data gathered by the SPARKL workflow can reveal deeper insights regarding the mechanism of cell death when utilizing a dual-labeling approach. Cells dying by apoptosis sequentially label with Annexin V and then with a viability dye while cells dying by ferroptosis exhibit similar kinetics for both labels. In stark contrast, necroptotic cell death exhibited the reverse phenotype in which cells robustly label with the viability dye and then slowly accumulated Annexin V over time. It was recently reported that cells can exhibit Annexin V labeling prior to viability dye labeling due to the formation of PS-containing “bubbles” on the plasma membrane following MLKL activation (Gong et al., 2017). However, this phenotype was demonstrated to be rapid, transient, and required highly-sensitive technologies to detect. While we saw no evidence of this phenotype, we believe this is a consequence of the difference in signal detection sensitivity and time resolution between the methodologies and therefore do not believe the results to be inherently contradictory. Furthermore, these differences exemplify the importance of utilizing multiple specialized techniques to further understanding the molecular mechanisms of cell death.

### Kinetic data gathered by SPARKL provides several quantitative metrics for investigative and comparative analyses of cell death at both the single-cell and population level

Cell death assays using flow cytometry provide only endpoint data and are typically visualized as comparative bar graphs, quadrant plots of probe positivity, or histograms of fluorescent intensity. SPARKL is less invasive to cultured cells and is more data-rich due to both the collection of kinetic data and underlying cell imaging methodology. However, while the utilization of real-time live-cell imaging is becoming more commonplace, most users do not fully explore the depth of the technology or the data it generates when analyzing their experiments. Furthermore, the advantages of these technologies are further minimized by use of endpoint data, simplified comparisons, and labeling methods repurposed from flow cytometry (Goodall et al., 2016). Therefore, we aimed to broaden the landscape of experimental designs conducted with high-content imagers while interrogating how data generated by SPARKL could best be utilized and analyzed in order to examine phenomena currently overlooked by analogous workflows.

In the previous section, we characterized the labeling profiles for cells instigated to die by several cell death pathways as observed by real-time live-cell imagers. Not only did the labeling order differ between each pathway, but the kinetics and sensitivity of those labeling events were also observably different. Therefore, we do not suggest that either Annexin V or YOYO3 (or similar cell impermeable viability dye) is superior but rather the in tandem use provides new methods to understand the kinetics of cell death in response to a perturbagen. To demonstrate this, we investigated methods to quantify, process, and express the labeling profiles for each cell death pathway. To articulate the relative labeling kinetics for each pathway, we expressed kinetic data as the difference between Annexin V- and YOYO3-positive events over time (Figure 3A). Representation of data in this manner clearly identifies the period when differential labeling is most observed and how this period is variable between treatments or pathways of cell death. Interestingly, applying the same method to data collected from WT and KO AEFs revealed substantially dissimilar sequential labeling, also in the TNFα+Brp alone treatment (Figure 3B). Thus, these data suggest that TNFα+Brp-induced cell death in KO AEFs was more apoptosis-like but signaled by a different mechanism in the WT AEFs. Inspecting the labeling data taken from a coculture of WT and KO AEFs clearly demonstrated how labeling phenotypes are more difficult to identify in heterogeneous populations. The cocultured AEFs retained similar labeling kinetics of Annexin V but exhibited notably different YOYO3 labeling when compared to either homogeneous population, such as the separation of the zVAD+Nec1 co-treatment (Figures 3C and 2G). When we compared the labeling data by expressing data as the difference between relative Annexin V- and YOYO3-positive events, we observed a trend concomitant with the combination of the two cellular populations (Figures 3D).

**Figure 3.**
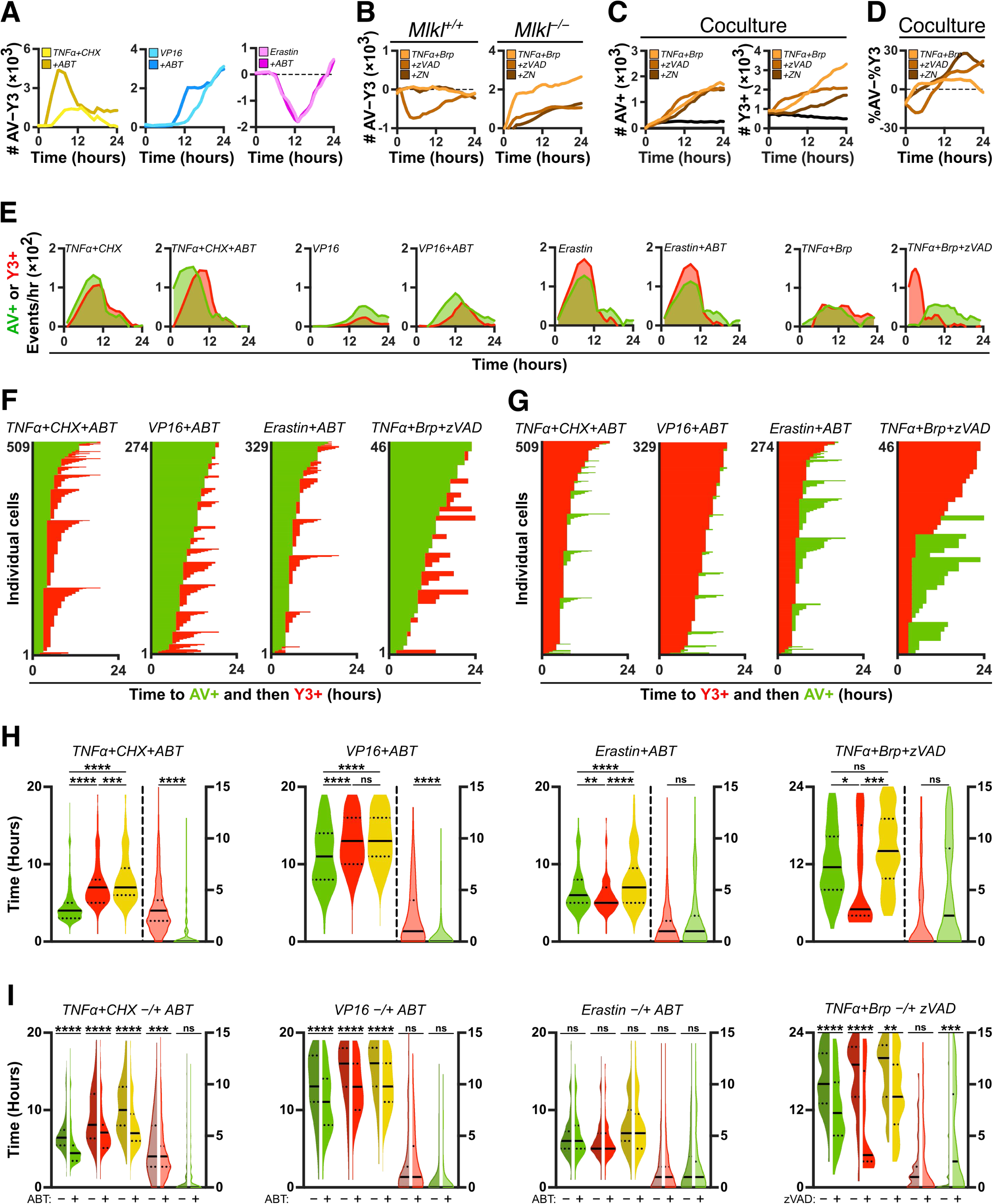
A dual-reporter SPARKL workflow for characterizing modes of cell death at population and single-cell level. (**A**) Number of YOYO3-positive events was subtracted from the number of Annexin V-positive events for cells analyzed in ***Figure 2B***. (**B**) Number of YOYO3-positive events was subtracted from the number of Annexin V-positive events for cells analyzed in ***Figure 2G***. (**C**) WT and *Mlkl* KO AEFs were cocultured in a 1:1 ratio, treated with mTNFα (50 ng/ml) + Birinapant (10 µM), and co-treated with zVAD-fmk (40 µM), with or without necrostatin-1 (10 µM), and subjected to SPARKL in the presence of Annexin V-FITC (250 ng/ml) and YOYO3 (250 nM). (**D**) Annexin V- and YOYO3-positive events for cells treated and analyzed in ***C*** were normalized as a percent of total signal and the percent YOYO3-positive was subtracted from the percent Annexin V-positive for each time point. (**E**) Histogram data from cells treated and analyzed as in ***A-D*** shown as number of new Annexin V (green) or YOYO3 (red) events occurring during each hour. (**F**) WT MEFs were treated with CHX (50 μg/ml), mTNFα (10 ng/ml) + CHX (50 μg/ml), VP16 (25 μM), or Erastin (10 μM), and co-treated with ABT-737 (1 μM); WT AEFs were treated with mTNFα (50 ng/ml) + Birinapant (10 µM) and co-treated with zVAD-fmk (40 µM). Cells were subjected to SPARKL in the presence of Annexin V-FITC (250 ng/ml) and YOYO3 (250 nM). Individual cells within the population imaged by the machine were tracked and analyzed to identify the time at which a cell was single- and double-positive. Panels show analysis where single-positive is defined as Annexin V-positive and double-positive is defined by subsequent labeling with YOYO3. (**G**) Same data as in ***F*** but panels show analysis where single-positive is defined as YOYO3-positive and double-positive is defined by subsequent labeling with Annexin V. (**H**) WT MEFs and WT AEFs treated and analyzed in ***F***. Population distribution of time-to-signal is shown for the following metrics: time to Annexin V (green), time to YOYO3 (red), time to double-positive (yellow), time to YOYO3 after Annexin V (transparent red), and time to Annexin V after YOYO3 (transparent green). Green, red, and yellow plots are scaled to the left axis while the transparent plots are scaled to the right axis. (**I**) WT MEFs and WT AEFs were treated and analyzed as in ***F*** and ***G*** without co-treatment of ABT-737 or zVAD-fmk, respectively. Violin plots of collected data were compared with data from ***H*** in which cells were co-treated as indicated. Green, red, and yellow plots are scaled to the left axis while the transparent plots are scaled to the right axis. Error bars are not shown for kinetic data to aid in data visualization. Violin plots display the entire population of collected data and have been smoothened to aid in visualization Instances of flat ends are an artifact of the method used to create the frequency distribution and do not reflect missing or truncated data points. Median and quartiles are shown as a solid and dotted line, respectively. Population statistics were conducted by performing a Mann-Whitney *U* test for each indicated comparison. Significance is noted as follows: \*\*\*\**P* < 0.0001, \*\*\**P* < 0.001, \*\**P* < 0.01, \**P* < 0.05, ns = not significant.

It is important to note that the kinetic data collected by SPARKL represents populations of dying cells labeling with fluorescent probes. For an individual cell, detection of fluorescent labeling is a binary process and pixels are considered “positive” when they exhibit fluorescence above a threshold value established during algorithm training (see Materials and Methods). We utilized the AEF genetic model as an exaggerated example to illustrate how data interpretation can be less clear in heterogeneous populations. However, heterogeneity in response to a perturbagen, and therefore the kinetics of cell death and labeling, is the culmination of many biological and stochastic factors and is not necessarily so overt in isogenic populations (Roux et al., 2015; Inde and Dixon, 2018). Therefore, we explored methods in which SPARKL could assess heterogeneity in a population dying in response to a perturbagen. Instead of viewing data as cumulative, kinetic data can be processed and expressed as the number of new events over time. Comparing histograms of events per hour for both labels revealed a more rapid labeling of Annexin V compared to YOYO3 in apoptotic models, overlapping histograms in ferroptosis, and stark YOYO3-first sequential labeling in cells undergoing necroptosis (Figure 3E). These data better illustrate labeling trends within the cell population and therefore can be analyzed for population statistics (such as reporting mean, median, and deviation of cell events within the population). Additionally, data expressed in this manner can reveal subpopulations that label differentially and therefore could reflect different biological mechanisms. For instance, we consistently observed “shelves” which were most noticeable when the perturbagen resulted in rapid and robust cell death, such as either Annexin V or YOYO3 for TNFα+CHX or YOYO3 for TNFα+Brp+zVAD.

High-content live-cell imagers observe the same cells over time and provide the opportunity for investigators to track labeling kinetics on a single-cell level. We utilized this technological advantage and applied it to our dual-labeling workflow in order to assess and quantify sequential labeling for each cell within the imaging field. Image stacks and analysis masks generated during the SPARKL workflow were processed using ImageJ and Fiji (Schneider et al., 2012, Schindelin et al., 2012). Fluorescent events were spatially and temporally defined as regions of interest (ROIs) and then cross-referenced between images slices and fluorescent channels. Analyzed in this way, the time-to-signal for both fluorophores was determined for each observed cell. WT MEFs and AEFs were instigated to die, analyzed by this method, and graphed as the time to single-positive followed by time to double-positive. Interestingly, while labeling kinetics were treatment-specific, the time in which Annexin V-labeled cells sequentially labeled with YOYO3 was highly variable (Figure 3F). For example, 75% of MEFs treated with TNFα+CHX+ABT labeled with Annexin V by 5 hours and co-labeling with YOYO3 occurred, on average, 3.2 hours later but the slowest 10% were highly variable. This analysis observed cells once they became Annexin V-positive and then assessed the time in which the cell became double positive. As a consequence, ROIs which labeled either simultaneously or in reverse order are visualized as a bar with no red phase and cells exhibiting this pattern appeared more frequently and rapidly in models of non-apoptotic cell death. To corroborate this observation, the same image stacks were reanalyzed to track the time to YOYO3- and then double-positivity (Figure 3G). Very few cells instigated to undergo apoptosis exhibited Annexin V labeling after YOYO3 (visualized by the lack of green phase in most of the bars) but ferroptotic cells exhibited approximately 50% of YOYO3-first sequential labeling. Interestingly, this analysis revealed a dramatic bifurcation within the population of treated AEFs: while half of the population demonstrated a phenotype consistent with necroptosis (i.e., rapid onset of YOYO3 followed by variable Annexin V labeling), the other half exhibited a phenotype reminiscent of a mitochondrial pathway of apoptosis (i.e., lengthy and highly variable labeling time, YOYO3 labeling after Annexin V). This phenotype could be a consequence of birinapant-mediated inhibition within the cell or the association of oligomeric MLKL with membranes other than the plasma membrane, such as the mitochondria (Wang et al., 2014).

SPARKL data analyzed and graphed in this manner aids in visualization of population trends by separately tracking the labeling kinetics of each individual cell. Therefore, labeling metrics of the population can be statistically compared to further characterize pathway- or treatment-specific patterns of cell death (Figure 3H). Sequential labeling can be confirmed and quantified by comparing individual label kinetics to double-positive kinetics. For example, YOYO3 and double-positive kinetics of MEFs treated with VP16+ABT are not significantly different which indicated that double-positivity depended on YOYO3 and that Annexin V labeling occurred first. Similarly, the reverse phenotype was observed in AEFs undergoing necroptosis even though the population exhibited two different labeling profiles. Interestingly, MEFs undergoing ferroptosis exhibited significantly distinct labeling kinetics for Annexin V, YOYO3, and double-positivity, which indicated no sequential labeling and likely reflects the biochemistry of the currently unknown method by which lipid peroxidation results in cell death (Feng and Stockwell, 2018). Additionally, this method for comparing cellular populations provides a convenient strategy to investigate the consequences of co-treatments and determine either synergistic or differential biology (Figure 3I). For example, MEFs induced to undergo apoptosis exhibited significantly different labeling kinetics in response to ABT-737 while ferroptotic MEFs did not. Importantly, the pattern of distributions, while occurring earlier, were conserved and therefore reflected a treatment-mediated synergy of the death pathway, namely apoptosis. In stark contrast, the switch from apoptosis to necroptosis in AEFs exhibited significantly different labeling kinetics as well as different patterns of distribution. Collectively, these data demonstrate a method and rationale for analyzing data collected by high-content live-cell imagers to characterize and compare differential labeling kinetics of single cells in response to perturbagens.

While there are instances to do the deep analyses we described above, it does not represent the common usage of high-content live-cell imagers for cell death analyses – high-volume comparative studies. In order to take full advantage of the high-throughput capacity of these technologies, complex kinetic data must to be expressed as simplified key parameters. Here, we will review best practices for comparing SPARKL data as well as several mathematical methods for rapid parameterization of collected data (Figure 4A). Instinctually, investigators often use the maximum number of events to compare cell death data and use language akin to “A had more or less death than B.” However, the amplitude of kinetic curves reflects extraneous factors such as the number of cells in the population or the penetrance of a particular perturbagen. This is why graph axes are set by control treatments and do not need to be the same scale between cell types (such as data in Figure 1E). Additionally, using the maximal number of events (i.e.: death) is highly reductionist and does not reflect metrics that are uniquely collected by real-time methodologies such as the times of onset, 50% death, plateau, maximal rate of death, etc. – collectively referred to as the “kinetics” of cell death. Therefore, we recommend drawing comparisons from cell death kinetics.

**Figure 4.**
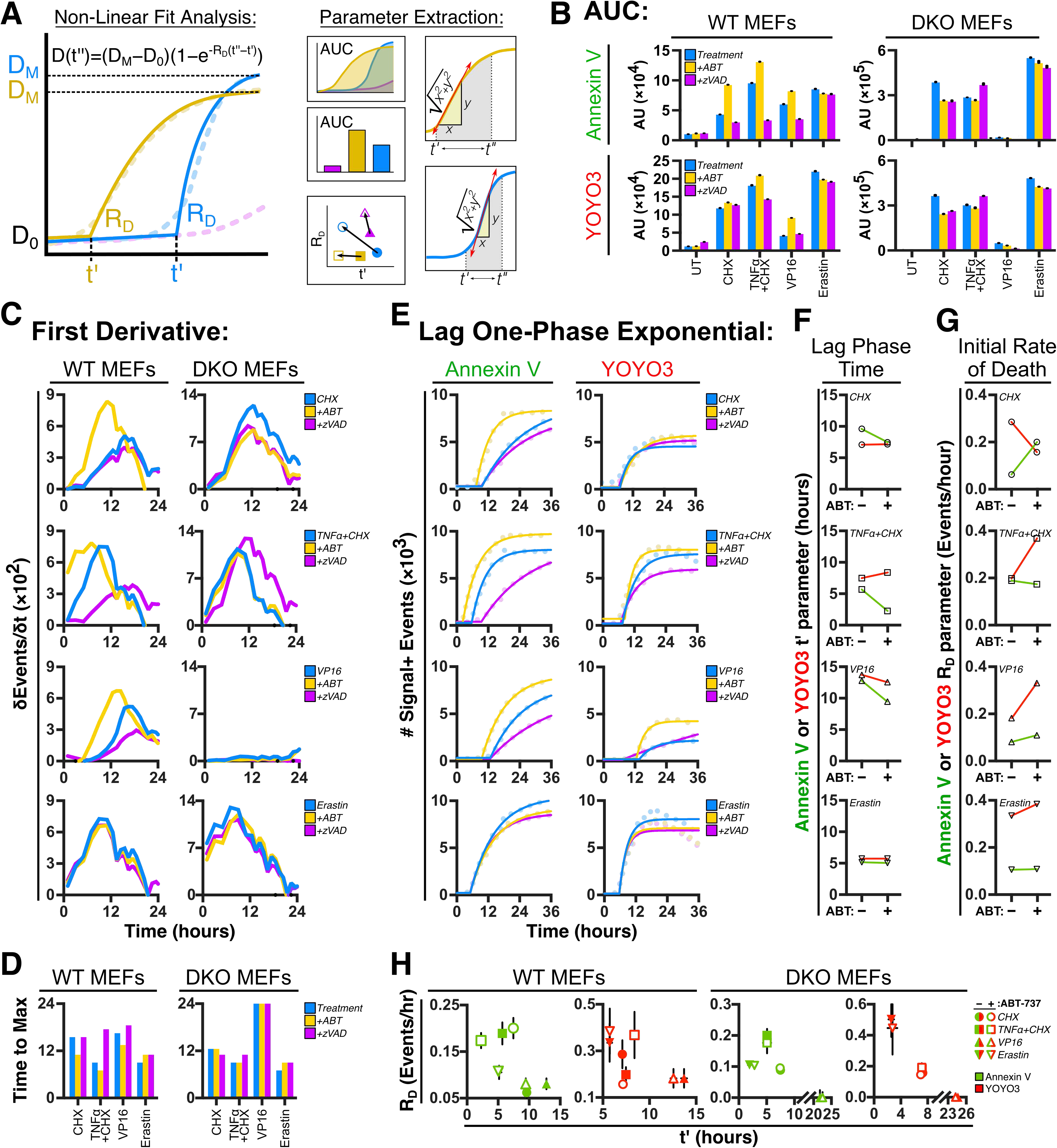
Mathematical analysis of the SPARKL kinetic data provides several parameters for high-throughput comparative assays. (**A**) Example methods for mathematically analyzing kinetic data and extracting parameters for cell death comparative studies. Area under the curve (AUC) provides a single parameter for comparing kinetic data between treatments or cell types. Mathematical conversion of kinetic data can provide additional comparative parameters such as death rates via derivatization or death onset from non-linear fit analyses. (**B**) WT and DKO MEFs were incubated in DMEM containing Annexin V-FITC (250 ng/ml) and YOYO3 (250 nM), treated with CHX (50 μg/ml), mTNFα (10 ng/ml) + CHX (50 μg/ml), VP16 (25 μM), or Erastin (10 μM), co-treated with DMSO, ABT-737 (1 μM), or zVAD-fmk (100 μM), and subjected to SPARKL. Area under the curve (AUC) values were calculated from kinetics of Annexin V- and YOYO3-labeling. Dissimilar range of axis reflects differences in number of total cells due to rates of proliferation. Axis range is determined by internal controls within each experiment. (**C**) First derivative of data collected from cells treated as in ***B*** depicting the rates of population cell death at each time point. (**D**) Comparison of maximum rates of death as calculated from first derivatives in ***C***. (**E**) Fit curves using a Lag One-Phase Growth function (solid line) applied to data from cells treated in ***B*** (dotted line). Curve for YOYO3 labeling kinetics in cells treated with VP16 and zVAD-fmk was considered an ambiguous fit and therefore excluded from parameter extraction and comparisons. (**F**) Comparison of lag phase time (*t’*) from Annexin V and YOYO3 data fits for cells treated as indicated from ***E***. (**G**) Comparison of initial/maximal rate of death (*RD*) from Annexin V and YOYO3 data fits for cells treated as indicated from ***E***. (**H**) 2D graphs of parameters calculated in ***E*** and graphed in ***F*** and ***G*** provide a convenient method for comparing treatment trends across labels and cell types. Error bars are not shown in instances where they are smaller than the data symbol.

The simplest method is to calculate the area under the curve (AUC) which compresses kinetic data into a single parameter for convenient comparisons. WT and DKO MEFs were instigated to die using a variety of perturbagens, co-treated with ABT-737 or zVAD-fmk, and detected by Annexin V- and YOYO3-labling (Figure 4B). AUC values were greater in cells co-treated with ABT-737 and reduced in cells co-treated with zVAD-fmk for perturbagens activating apoptosis, but unchanged in cells undergoing ferroptosis. AUC calculated from YOYO3 detection exhibit similar trends in response to ABT-737 and zVAD-fmk albeit to a lesser degree (Figure 4B). These analyses retained the observed trends of Annexin V and YOYO3 labeling within the raw kinetic data. Alternative metrics of interest are the rate of cell death and the time to maximal rate of death. The first derivative was calculated for data collected from MEFs treated with pro-death perturbagens and labeled with Annexin V (Figure 4C). The derivative reports the rate of cell death within the population, or the slope of kinetic curves, at each collected time point. MEFs undergoing apoptosis demonstrated greater rates of cell death at earlier time points in response to ABT-737 while exhibiting lower rates in response to zVAD-fmk (Figure 4C, yellow and purple data). Additionally, data can be presented as the time-to-maximal rate of death, however this parameter is only informative for samples exhibiting significant death and labeling over the course of data collection (Figure 4D). These mathematical analyses represent simple but robust methods to parameterize data collected with SPARKL for convenient comparisons between experimental conditions.

In an effort to quantify data in a manner that better retains the richness of kinetic data, we employed a non-linear fit analysis to the data collected with SPARKL. A Lag One-Phase Exponential (LOPE) function provided the best fit for our data and has been utilized successfully by other groups for modeling cell death kinetics (Forcina et al., 2017) (Figure 4E). This mathematical function reflects the latency period prior to the onset of cell death in which cells undergo biological signal transduction in response to perturbagens. The LOPE function calculates two useful parameters during the fit analysis: the duration of the lag phase (*t’*) and the rate of cell death following the lag phase (*RD*). It is worth noting that this model assumes that *RD* is the maximal rate of cell death which is not always accurate. However, this model consistently performed better when fitting data compared to other models (for example, sigmoidal functions which required more defined asymptotes to model). Similar to graphs of raw kinetic data, the plateau amplitude is dependent on the number of cells and therefore was not considered for parameterization in comparative studies. Comparing *t’* values for WT MEFs labeled with Annexin V and undergoing apoptosis demonstrated a clear trend in which ABT-737 co-treatment reduces *t’*, while the *t’* of MEFs undergoing ferroptosis was unchanged (Figure 4F). *RD* trends calculated from Annexin V data were harder to interpret since the metric only differed in MEFs co-treated with CHX and ABT-737 (Figure 4G). We believe this may be an indication that the selected perturbagen concentrations were excessive and therefore *RD* has plateaued, or an indication that the time of maximal death significantly differs from the *RD* calculated using LOPE. Interestingly, *t’* from YOYO3 data did not reflect changes in response to ABT-737 and *RD* values increased (Figures 4F-G, red data). Viewing these data as two-parameter plots reveals treatment- or cell type-specific trends in *t’* and *RD* following co-treatment with ABT-737 (Figure 4H). This strategy for data visualization is particularly effective when characterizing cell death mechanisms for perturbagens of unknown biology and conducive to large-scale, multi-parametric drug panels (Forcina et al., 2017; Fallahi-Sichani et al., 2013). While we have demonstrated methods that utilized both Annexin V and YOYO3, this method for comparative studies is most effective when utilizing a single labeling agent, whichever is most relevant to the experimental design.

The advent of high-content live-cell imagers provides investigators the opportunity to collect and analyze cellular data in a high-volume, high-throughput manner in real time. However, these technologies are not often used to their fullest potential and instead serve as a surrogate for flow cytometric workflows. To capitalize on the advantages of these technologies, we have provided several methods and best practices for utilizing and interpreting the data in comparative studies. We described a multiplex labeling workflow to demonstrate the variety of experimental designs compatible with these machines as well as several methods to parameterize and appropriately compare collected data. Collectively, these examples showcase the versatility, range of applications, breadth and depth of collected data, and analyses investigators can apply to experimental paradigms by using the SPARKL workflow.

### Simultaneous collection of proliferation and cell death data to accurately characterize perturbagen-induced cytostatic and cytotoxic effects

The high-throughput capacity of SPARKL permits investigators to conduct exhaustive optimizations and drug titrations with relative ease. However, perturbagens at sub-lethal concentrations may still affect rates of cell proliferation, either directly or as a consequence while mitigating the stress signal. Additionally, it has been shown that growth rate inhibition metrics can correct otherwise confounding data in large-scale drug screens (Hafner et al., 2016). Therefore, investigators require a method to properly assess the cytostatic effects of perturbagens on cells in culture. This is commonly accomplished by either measuring confluency or surface area metrics, which do not accurately reflect cell numbers, or by requiring the investigator to generate cell lines expressing fluorescent proteins, which is laborious, limiting to model systems, and not conducive to high-throughput analyses (Artymovich and Appledorn, 2015). Previously, we described a method utilizing non-toxic CFSE to label cells prior to treatment and data acquisition (Gelles and Chipuk, 2016). This method required additional handling steps, suffered mild photobleaching, and was not compatible with certain cell types. Therefore, we sought to develop a label-and-go protocol of SPARKL that would simultaneously collect proliferation and cell death data in real-time without requiring a separate cell-labeling step (Figure 5A).

**Figure 5.**
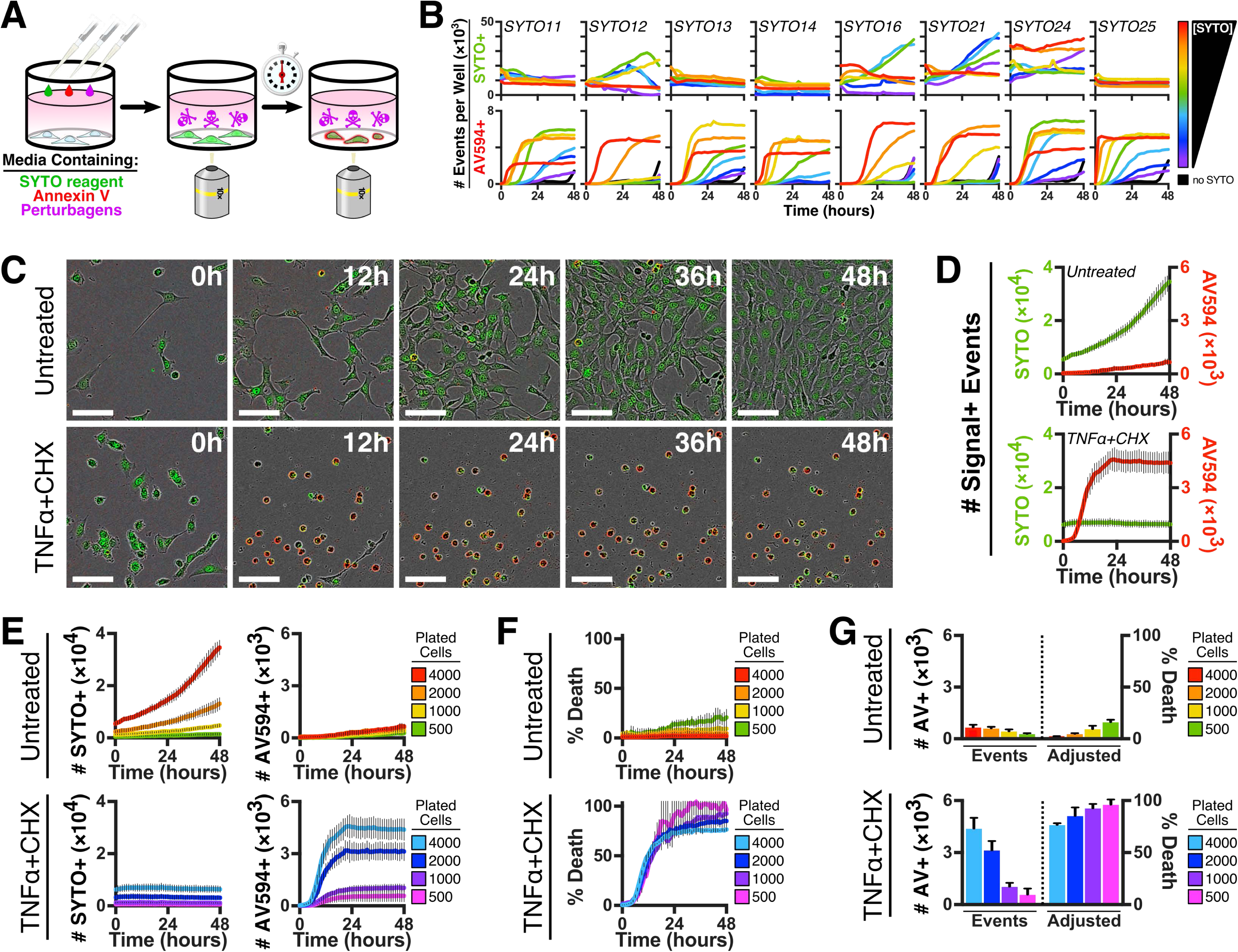
Non-toxic label-and-go reagents used with SPARKL can normalize data to account for changes in cell proliferation. (**A**) Label-and-go method to collect proliferation data during Annexin V-binding assay for composite cytostatic and cytotoxic effects of perturbagens. (**B**) WT MEFs were incubated with titrations of SYTO nucleotide labeling reagents (0.03-20 μM) in the presence of Annexin V-594 (1 μg/ml) and subjected to SPARKL in order to assess toxicity of the SYTO dyes. (**C**) WT MEFs were co-treated with CHX (50 μg/ml) and mTNFα (10 ng/ml) and incubated with Annexin V-594 (1 μg/ml) and SYTO21 (300 nM). Images captured at indicated time-points. (**D**) Kinetic data analyzed from images stacks corresponding to cells from ***C***. (**E**) WT MEFs were seeded at the indicated numbers 24 hours prior to treatment with mTNFα (10 ng/ml) + CHX (50 μg/ml) and incubation with Annexin V-594 (1 μg/ml) and SYTO21 (300 nM). Cells were then subjected to SPARKL to analyze proliferation and cell death kinetics. (**F**) Cell death data from ***E*** was normalized using SYTO21 and expressed as percentage of the population exhibiting death (% Death). (**G**) Endpoint data of Annexin V-594-positive events from ***E*** and corresponding percent death (% Death) normalization from ***F***.

Probes labeling the nucleus provide robust and readily detectable fluorescent events, which can be efficiently analyzed for accurate quantification of event number within each sample. However, most nuclear labeling probes are either cell impermeable or function as toxic DNA intercalators which are incompatible with time-course experiments and the SPARKL workflow. Therefore, we investigated if cell-permeable, low-affinity cyanine nucleic acid stains were suitable for prolonged incubation with cells. MEFs were incubated with varying concentrations of SYTO fluorescent dyes and Annexin V conjugated with AlexaFluor 594 (Annexin V-594) in order to assess the cytotoxic effects of nucleotide labeling (Figure 5B). Each tested SYTO dyes exhibited toxicity at high concentrations and degrees of cytostatic behavior at low- to mid-range concentrations. Low-concentration ranges of SYTO16 and SYTO21 demonstrated no adverse effects to cell proliferation or viability, and we ultimately selected SYTO21 for use within SPARKL workflows. SYTO21 primarily labeled cells in the nucleus, was readily detected and segmented, and capable of labeling daughter cells post-mitosis (Figure 5C). Additionally, SYTO21 signal was retained in cells undergoing apoptosis and resulted in SYTO21/Annexin V-594 double-positive events. MEFs incubated with SYTO21 demonstrated an increase in SYTO21-positive events over time, indicative of proliferation. By contrast, MEFs undergoing apoptosis via treatment of TNFα+CHX exhibit no change in the number of SYTO21-positive events and robustly labeled with Annexin V-594 (Figure 5D). Therefore, SYTO21 is suitable as a fluorescent probe to track cell proliferation and population growth.

High-content live-cell imagers detect fluorescence and express data as a number of positive events within a given area. However, this value is not normalized for total cell number and therefore more events are detected in wells containing more cells. Interpretations of data expressed in this manner may be obfuscated when cell numbers vary between samples due to either treatment-induced or cell type-specific differences in proliferation rates. Imperfect strategies to circumvent this issue undermine the advantages of kinetic monitoring by either implementing tandem analysis via flow cytometry, normalizing to end-point data collected using DNA-intercalating dyes or potent inducers of cell death, or using confluency as a surrogate for cell number (Lopez et al., 2016; Giampazolias et al., 2017). Therefore, we investigated whether SYTO21 labeling could be used to normalize cell death data collected with SPARKL. MEFs were seeded at several densities, incubated with SYTO21 and Annexin V-594, and treated with either vehicle or TNFα+CHX. SYTO21 data accurately measured differences in cell number and tracked cell proliferation in untreated MEFs; MEFs instigated to die exhibited constant SYTO21 data and the concomitant Annexin V-positive data exemplified how cell number can complicate data interpretation (Figure 5E). The number of Annexin V-positive events at each timepoint was normalized by the corresponding SYTO21-positive events to express data as the percent of the population exhibiting Annexin V-positivity (Figure 5F). For comparison, Annexin V-positive endpoint data collected using SPARKL is shown both as the raw number of events and the adjusted percentages normalized using total cell number as reported by SYTO21 (Figure 5G). These data demonstrate that when coupled with skilled cell culture practices, this dual-labeling method and analysis can control for cell number differences due to differential cell seeding and cell type-inherent or treatment-induced alterations to proliferation rates.

Cell death data normalization is particularly relevant when perturbagens do not illicit a rapid death phenotype, which will result in more proliferation variability between experimental conditions and therefore different maximal labeling. To demonstrate this, we applied the normalization methodology to WT and DKO MEFs, which exhibit dramatic differences in rates of proliferation, incubated with SYTO21 and Annexin V, and treated with a panel of perturbagens. Annexin V data collected through SPARKL was normalized by the SYTO21 data and provided values that can be directly compared between the two cell types (Figure 6A). Perturbagens engaging BAX- and BAK-mediated apoptosis demonstrated an attenuated lethal response in DKO MEFs (such as VP16 and calcium ionophore, A23187) and perturbagens capable of engaging the intrinsic pathway of apoptosis were synergized by co-treatment with ABT-737. When we replicated this experiment using YOYO3 as the cell death probe and normalized for SYTO21 labeled cells, we observed a significant loss of signal during longer experiments (Figure 6B). When analyzed for longer than 24 hours, MEFs undergoing apoptosis by co-treatment with TNFα+CHX exhibited sustained SYTO21- and Annexin V-positivity, but demonstrated a loss of YOYO3 signal over time (Figure 6C). We analyzed the average fluorescent intensity of positive events which revealed that YOYO3-positive events lost fluorescence over time (Figure 6D). While photobleaching may contribute to this loss of signal, we believe that the effect is indicative of changes to DNA content and subsequent labeling by YOYO3, which would explain why the diminished signal is perturbagen-specific and observed in cells undergoing apoptosis. Signal loss does not appear to be due to cells leaving the focal plane as this would affect detection of all probes and the data do not demonstrate this concomitant loss of signal in either SYTO21 or Annexin V over time.

**Figure 6.**
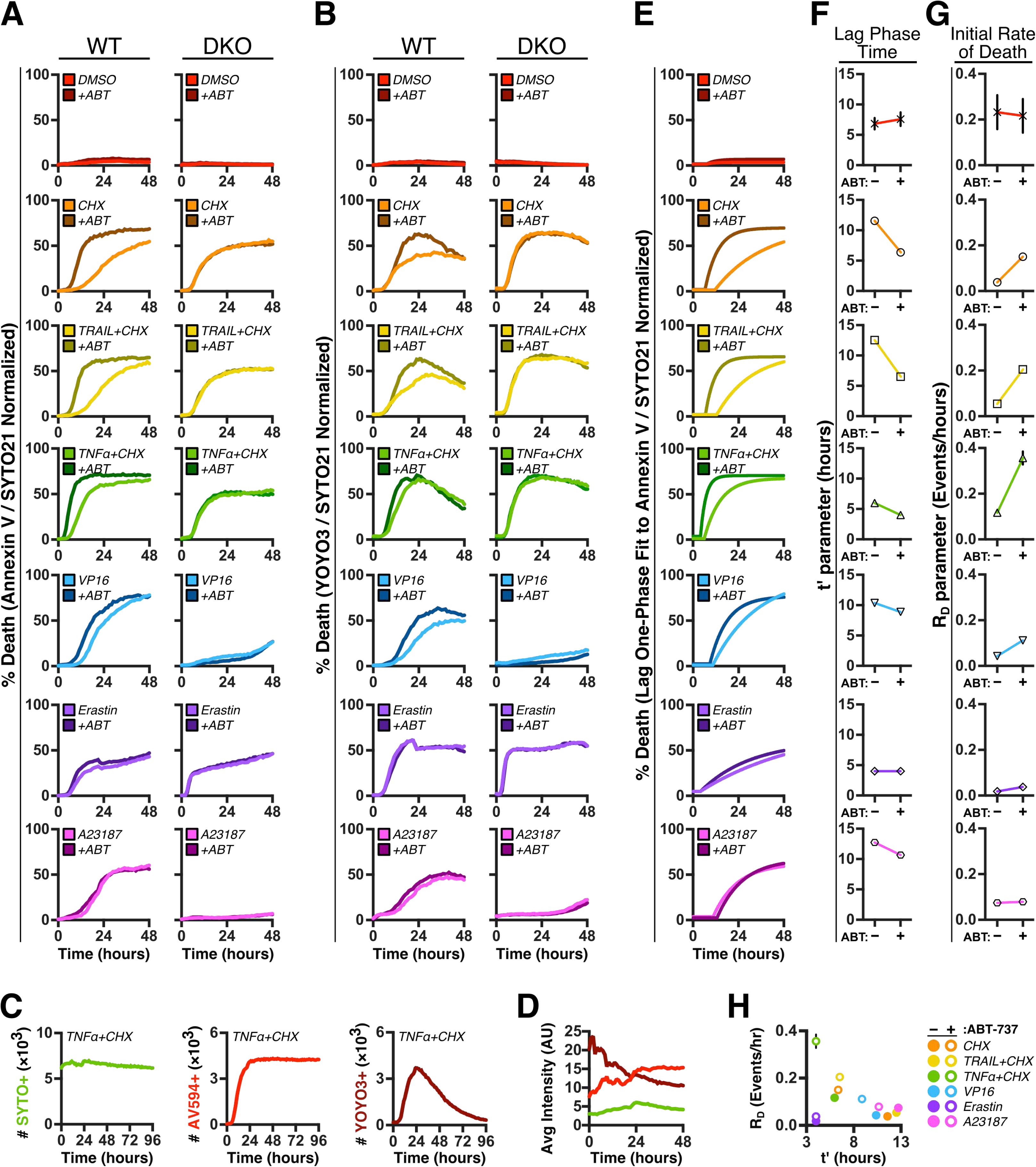
Parameters extracted from non-linear fit analyses of normalized death kinetics generate comparative profiles for studying mechanisms of cell death. (**A**) WT and DKO MEFs were incubated in DMEM containing SYTO21 (300 nM), Annexin V-549 (1 μg/ml), indicated treatments of CHX (50 μg/ml), TRAIL (25 ng/ml) + CHX (50 μg/ml), mTNFα (10 ng/ml) + CHX (50 μg/ml), VP16 (25 μM), Erastin (10 μM), or A23187 (5 μM), co-treated with either DMSO control or ABT-737 (1 μM), and subjected to SPARKL workflow. Percent death (% Death) was calculated by Annexin V data normalized by the SYTO21 data at each timepoint. (**B**) Cells, treatments, and analysis as in ***A*** for cells cultured in DMEM containing SYTO21 (300 nM) and YOYO3 (250 nM). (**C**) WT MEFs treated as in ***A*** were subjected to long-term, repetitive detection of SYTO21-, Annexin V-594-, and YOYO3-positive events to assess signal permanence and longevity. (**D**) Average integrated intensity of events detected in ***C*** showing changes in fluorescent intensity of reporters over the first 48 hours. (**E**) Fit curves using a Lag One-Phase Growth function applied to data from WT cells in ***A***. Curve for DMSO-treated cells was considered an ambiguous fit and therefore its parameterization is variable. (**F**) Comparison of lag phase time (*t’*) from fits applied to cells treated as indicated in ***A***. (**G**) Comparison of initial/maximal rate of death (*RD*) from fits applied to cells treated as indicated in ***A***. (**H**) 2D graph of parameters calculated in ***E*** and graphed in ***F*** and ***G*** provides a convenient method for comparing trends across various cytotoxic treatments. Error bars are not shown for kinetic data to aid in data visualization, and not shown in instances where they are smaller than the data symbol.

We capitalized on this method to normalize data and applied the same mathematical analyses described in the previous section. Percentages of cell death calculated by normalizing Annexin V data to SYTO21 were fit using the LOPE function (Figure 6E). Samples co-treated with ABT-737 reveal a reduction in lag phase time (*t’*), indicative of BCL-2 family contributions within the cell death pathway; by contrast, *t’* is unchanged in response to ABT-737 in MEFs undergoing ferroptosis via erastin (Figure 6F). Consistent with our previous methods of analysis, receptor-mediated apoptosis by either TNFα or TRAIL in conjunction with CHX demonstrated substantially different kinetics when co-treated with ABT-737 and indicates a predominance for Type II signaling in MEFs. When analyzing the initial rates of cell death (*RD*), which became more informative for data which had been cell number-normalized, we observed increased rates in of death following co-treatment of ABT-737 in MEFs treated with CHX, TNFα+CHX, TRAIL+CHX, or VP16 (Figure 6G). *RD* was unchanged in MEFs treated with erastin or A23187. Cell death data from MEFs treated with A23187 collectively indicates a mechanism requiring BAX and BAK but only partially sensitized by inhibition of anti-apoptotic BCL-2 proteins, which is a profile not shared amongst the other perturbagens exampled within this work (Figure 6H). Control-treated MEFs demonstrated ambiguous fit curves and therefore parameter extraction was highly variable and not informative (Figures 6E-G, red panel). Similarly, DKO MEFs not exhibiting death in response to particular perturbagens (such as VP16 and A23187) produced ambiguous fit curves and were not analyzed further (data not shown). These data exemplify the utility of mathematical parameterization on kinetic data which was normalized for differential cell numbers as a consequence of cell-inherent or treatment-induced changes to proliferation.

## Discussion

Here we described the SPARKL workflow which is capable of simultaneously gathering proliferation and cell death data at single-cell and population-level resolutions. Using multiplex fluorescent detection of cell death, investigators can dichotomize cells engaging apoptotic or non-apoptotic signaling pathways in response to perturbagens. Additionally, we described a novel method to quantify cell numbers and proliferation using a non-disruptive fluorescent probe which is compatible with “label-and-go” workflows conducive to high-throughput studies. Collection of these rich datasets provides in depth comprehensive insight into the cytostatic effects of perturbagens as well as their cytotoxic potential. Furthermore, we described and advanced robust and thorough mathematical analyses that parameterize data for convenient comparative analyses and interpretations of underlying cell death biology. The presented data demonstrate the versatility of the technology using single concentration treatments, but these workflows and analyses are well suited for experiments using perturbagen titrations and determining optimal conditions for subsequent studies. Collectively, SPARKL is a versatile, data-rich, non-intensive, and zero-handling method to visualize, detect, and quantify the kinetics of cell death which, when coupled with mathematical analyses, enable investigators to conduct thorough high-throughput comparative studies.

### Limitations

The data presented herein have utilized either immortalized MEFs or primary AEFs to demonstrate the workflows compatible with SPARKL. These adherent cells demonstrate minimal PS exposure under normal growth conditions. In our method, cells are incubated with labeled Annexin V which may bind stochastically-exposed PS on the outer leaflet of the plasma membrane and result in fluorescent puncta. These puncta are easily excluded while developing appropriate processing definitions, but may pose a difficulty for cell types demonstrating significant basal PS localization on the outer leaflet. Additionally, while MEFs tolerated the use of SYTO21 as a non-perturbing cell marker, other cell lines may demonstrate increased sensitivity to the nucleotide label, varying its efficacy. During analysis, accurate quantification of fluorescent events requires segmenting neighboring fluorescent signals without inappropriately bifurcating individual events. Therefore, cell lines that grow in colonies or exhibit a “cobble-stone” monolayer may require more stringent optimization to adequately establish segmentation between neighboring cells. Of note, apoptotic bodies demonstrated minimal detachment from the plate following cell death and labeled sufficiently with our detection reagents. However, alternative cell models or perturbagens may not remain adherent following cell death or may result in complex morphology that could complicate the quantification or detection of fluorescent events. While many of these applications can be modified for suspension cells adhered to the plate by a binding agent, additional troubleshooting is necessary and event segmentation is generally less representative due to cellular aggregates. Finally, high-content live-cell imagers require more robust labeling for detection when compared to flow cytometry and therefore certain methods, such as transient expression of fluorescent proteins, may not be detectable in these high-throughput workflows.

## Acknowledgements

This work was supported by: NIH grants CA157740 (J.E.C.), CA206005 (J.E.C.), AI52417 (A.T.T.), and F31AA024681 (J.D.G.), the JJR Foundation, the William A. Spivak Fund, the Fridolin Charitable Trust, an American Cancer Society Research Scholar Award, a Leukemia & Lymphoma Society Career Development Award, and an Irma T. Hirschl/Monique WeillCaulier Trust Research Award. This work was also supported in part by two research grants (5FY1174 and 1FY13416) from the March of Dimes Foundation, and the Developmental Research Pilot Project Program within the Department of Oncological Sciences at the Icahn School of Medicine at Mount Sinai, and the Tisch Cancer Institute Cancer Center Support Grant (P30 CA196521). *Mlkl^-/-^* mice were a gift to Adrian Ting from Warren Alexander at the University of Melbourne via Douglas Green at St. Jude Children’s Research Hospital.

## Materials and Methods

### Reagents and equipment

Cell culture reagents and chemicals were from Sigma Aldrich (St. Louis, MO, USA) or Thermo Fisher Scientific (Waltham, MA, USA) unless otherwise stated. Drugs and biologics as follows: ABT-737 (Abbott Pharmaceuticals, Lake Bluff, IL, USA); CHX (Sigma); YOYO3 and SYTO reagents (Life Technologies/Thermo Fisher Scientific), mTNFα and mTRAIL (Peprotech, Rocky Hill, NJ, USA); zVAD-fmk (ApexBio, Houston, TX, USA). Recombinant Annexin V was purified, labeled with FITC or Alexa Fluor 594 (Thermo Fisher Scientific), and re-purified using Dye Removal Columns (Thermo Fisher Scientific) following conjugation. Purity, stability, and fluorescence of labeled recombinant Annexin V was verified by SDS-PAGE and Flow Cytometry as previously described (Logue et al., 2009).

### Cell culture and treatments

Unless stated to the contrary, experiments were performed with *Bak^+/+^Bax^+/+^* and *Bak^−/−^Bax^−/−^* (WT and DKO, respectively) SV40-transformed MEFs (ATCC, Manassas, VA, USA). *Cyld^-/-^* MEFs were reconstituted with either a Flag tagged CYLD construct or empty vector as described previously (O’Donnell et al. 2011). Primary adult ear fibroblasts (AEFs) were isolated from *Mlkl^+/+^* and *Mlkl^-/-^* mice as follows: mouse ears were clipped, minced, and incubated in trypsin at 37°C for 1 hour with repeated vortexing before being cultured on tissue culture plates (Murphy et al., 2013). Cells were cultured in DMEM containing 10% FBS, 2 mM L-glutamine, and antibiotics and maintained in humidified incubators at 37°C with 5% CO2. For kinetic studies, cells were seeded at approximately 2 – 4 × 10^3^ cells/well in a 96-well format and left to adhere for 24 hours. Prior to live-cell imaging, growth media was replaced with complete phenol-red-free DMEM containing indicated fluorescent reagents and perturbagens. DMEM contains sufficient calcium concentrations for robust Annexin V labeling but other media formulations require calcium supplementation for optimal Annexin V labeling (1.5 – 2 mM CaCl2). Plates were pre-warmed prior to data acquisition to avoid condensation and expansion of the plate, which hinders auto-focusing during scan intervals. All tissue cultures were confirmed mycoplasma free with the colorimetric PlasmoTest detection kit (Invivogen, San Diego, CA, USA).

### SPARKL data acquisition and automated analysis

Kinetic experiments were performed with the IncuCyte ZOOM (Model 4459, Essen Bioscience, Ann Arbor, MI, USA) residing in a tissue culture incubator maintained as above, but other high-content live-cell fluorescent imagers are capable of replicating these data (e.g., IN CELL analyzer). Experiments were conducted for 24 – 96 hours with data collection every 1 – 2 hours to avoid photobleaching of fluorescent reporters. Using the 10× objective, a single plane of view was collected per well for 96-well plate assays. Phase contrast, green channel (Ex: 440/80 nm; Em: 504/44 nm; acquisition time: 400 ms), and red channel (Ex: 565/05 nm; Em: 625/05 nm; acquisition time: 800 ms) were collected for all experiments with spectral unmixing set as 3% of red removed from green. Images collected were 1392 × 1040 pixels at 1.22 μm/pixel. Automated image analysis was accomplished using the ZOOM software (V2018A) and defined by images collected using the specific cell lines and fluorescent reporters pertaining to the experiment. Processing definitions were designed using the settings for each fluorescent reporter as described below (Table 1). Data are expressed as events per well. The y-axis scale was determined for each experiment using internal controls to assess maximal death for a population. Fluorescent intensity data were analyzed using the average integrated intensity (AU × μm^2^) metric. Images of cell culture presented in this manuscript were white-point corrected for clarity and normalization of automated background subtraction in the green and red channel when no significant fluorescent event was detected. White-point correction was accomplished by adjusting temperature and tint parameters to overlap peak contributions of red, green, and blue pixels.

**Table 1:** Channel settings for each fluorescent reporter used for processing definitions with the IncuCyte ZOOM

|  | *Channel* | *Parameters* | *Radius*<br>*($\mu\text{m}$)* | *Threshold*<br>*(RFU)* | *Edge*<br>*Sensitivity* | *Area*<br>*($\mu\text{m}^2$)* |
| --- | --- | --- | --- | --- | --- | --- |
| Annexin V-AF594 | Red | Top-Hat | 10 | 5 | -35 | >150 |
| Annexin V-FITC | Green | Top-Hat | 25 | 3 | -30 | >100 |
| Annexin V-FITC (low) | Green | Top-Hat | 25 | 2 | -30 | >50 |
| SYTO21 | Green | Top-Hat | 25 | 0.5 | -25 | >150 |
| YOYO3 | Red | Top-Hat | 10 | 3 | -46 | >150 |

### Parameter extraction, mathematical processing, and data handling

Data exported from the IncuCyte ZOOM software were organized, handled, and analyzed using Excel v16.16.9 (Microsoft) and Prism 8.1.1 (Graphpad Software Inc.). Axis range for graphs was determined by internal negative and positive controls within each experiment. Event count data were converted into histograms using Equation 1, where #Signal^+^ refers to the number of events using either fluorescently-labeled Annexin V or YOYO3 for a given well “*i*” at time point “*j*”:

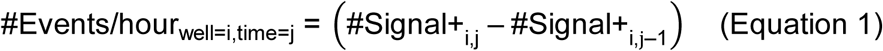

Area under the curve (AUC) values were calculated with the analysis tool in Prism and utilized the trapezoid rule ignoring peaks that were less than 10% of the distance from minimum to maximum Y values. The first derivative of collected data was calculated with the analysis tool in Prism and smoothed the data using 4 neighbors and 2nd order smoothing polynomial. Non-linear curve fits were calculated using the plateau followed by one-phase association function in Prism using the least squares fitting method given by Equation 2. Parameters extracted by Equation 2 are defined as follows: *D(t’’)* is the amount of death for given time point *t’’*; *D0* is the initial and minimum amount of death; *DM* is the plateau of maximal death; *RD* is the initial and maximal rate of death; *t’* is the time at which the lag plateau ends and exponential increase in death begins. All parameters were left unconstrained for mathematical fitting. Fits listed as ambiguous by Prism only occurred in conditions that did not result in death positivity and are highlighted in the text.

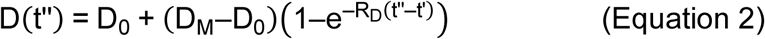

Experiments using SYTO21 to normalize data with cell counts was accomplished using Equation 3, where #Dead was determined by the event counts using either fluorescently-labeled Annexin V or YOYO3 for a given well “*i*” at time point “*j*”:

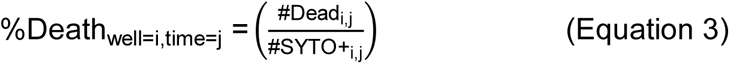

Unless otherwise noted, all experiments were conducted in triplicate and the presented data are the mean of replicates and representative of repeated experiments. When shown, error bars denote standard deviation of replicates; select graphs omit error bars for convenience of visualization.

### Single-cell two-hit kinetics

For experiments investigating sequential labeling of individual cells, analysis utilized event masks generated by the ZOOM software which were exported for further processing using external software packages. Specifically, masks for green and red channels as well as the overlap masks were used for these analyses. Cell event masks were recognized and spatio-temporally defined using ImageJ and Fiji software packages (ImageJ, National Institutes of Health, Bethesda, ML, USA) (Schneider et al., 2012, Schindelin et al., 2012). Images exported from the ZOOM software were imported into ImageJ as stacks for each mask collection (green, red, and overlay), converted to 8-bit grayscale and binary, and subjected to translation alignment using the StackReg and TurboReg plugins (Thévenaz et al., 1998). Reference regions of interest (ROIs) were autonomously defined by applying the Analyze Particles tool on the final image slice of the overlay mask. These reference ROIs were applied to all slices within the image stacks corresponding to the green, red, and overlay channels using the Multi-Measure tool. Positive pixel intensity within ROIs defined the slice in which the signal occurred. These data were then sorted and aligned in Excel by ROI reference number, thereby linking identified signal slices from each channel to the same ROI for graphing and interpretation. This workflow was used to quantify the progression of either green or red single-positivity to double-positivity for single cells within in a population.

